# Flux exponent control predicts metabolic dynamics from network structure

**DOI:** 10.1101/2023.03.23.533708

**Authors:** Fangzhou Xiao, Jing Shuang Li, John C. Doyle

## Abstract

Metabolic dynamics such as stability of steady states, oscillations, lags and growth arrests in stress responses are important for microbial communities in human health, ecology, and metabolic engineering. Yet it is hard to model due to sparse data available on trajectories of metabolic fluxes. For this reason, a constraint-based approach called flux control (e.g., flux balance analysis) was invented to split metabolic systems into known stoichiometry (plant) and unknown fluxes (controller), so that data can be incorporated as refined constraints, and optimization can be used to find behaviors in scenarios of interest. However, flux control can only capture steady state fluxes well, limiting its application to scenarios with days or slower timescales. To overcome this limitation and capture dynamic fluxes, this work proposes a novel constraint-based approach, flux exponent control (FEC). FEC uses a different plant-controller split between the activities of catalytic enzymes and their regulation through binding reactions. Since binding reactions effectively regulate fluxes’ exponents (from previous works), this yields the rule of FEC, that cells regulate fluxes’ exponents, not the fluxes themselves as in flux control. In FEC, dynamic regulations of metabolic systems are solutions to optimal control problems that are computationally solvable via model predictive control. Glycolysis, which is known to have minute-timescale oscillations, is used as an example to demonstrate FEC can capture metabolism dynamics from network structure. More generally, FEC brings metabolic dynamics to the realm of control system analysis and design.

## I. Introduction

Metabolism is the core interaction mechanism in biological systems across scales, from growth and survival in single strain microbial populations [1] to microbial communities [2] in human health [3], ecology [4], and metabolic engineering [5]. It is therefore important to study how the dynamics of metabolism is regulated. While the typical approach to study any dynamics in biological systems is through mechanistic models with many kinetic parameters identified via experiments (Fig. 1a), this is not feasible for dynamics of nontrivial metabolic networks. While we can experimentally measure bulk metabolic fluxes at scale and characterize metabolite stoichiometry robustly, we lack systematic ways to measure the kinetic parameters or observe the dynamic fluxes of intermediates in cells which depend on the concentrations of regulatory proteins and metabolites [6]. This data sparsity makes it impractical to identify the values of mechanistic parameters in nontrivial metabolic networks, let alone generalizing model predictions to situations different from the ones in fitted data.

**Fig. 1:**
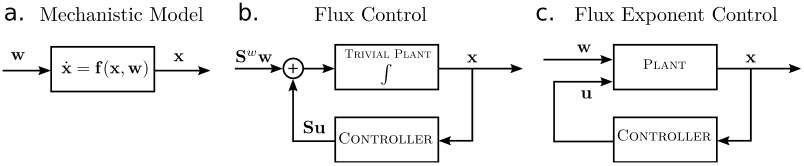
Control diagrams for several formulations of metabolic dynamics. **a**. The unstructured mechanistic description of metabolite concentration dynamics, with input as external exchange fluxes ***w*** and output as metabolite concentrations ***x*. b**. The flux control formulation, with stoichiometry ***S*** explicitly represented, and internal fluxes considered as control variable ***u***. The state variable ***x*** has trivial plant dynamics that is a direct integration of inputs. **c**. The flux exponent control formulation, with exponents of internal fluxes as control variable ***u***. The state variable ***x*** has nontrivial plant dynamics representing the uncontrolled internal fluxes.

In order to make progress despite sparse data, constraint-based approaches have been developed to model metabolism. Instead of trying to identify all the mechanisms and parameters in a system, a constraint-based approach takes known mechanisms as constraints and unknown mechanisms as free to vary, and looks at the set of all feasible behaviors (Fig. 3). *Flux control* is a constraint-based approach that identifies a natural known-unknown split between the stoichiometry and the flux of metabolic reactions, since the former is relatively easier to know while mechanisms of the latter is much harder to characterize. Therefore, flux control takes stoichiometry as a constraint on the dynamics of metabolite concentrations. This structure decomposes metabolism dynamics into a fixed stoichiometry and the changing fluxes. In other words, flux control splits the metabolic system into a plant-controller pair, with the stoichiometry as the plant and the fluxes as the controller, hence the name flux control (see Fig. 1b and 2). Then, either the set of all feasible fluxes can be analyzed for general rules, or optimization for certain objective functions such as growth maximization or ATP regeneration can be used to find specific points of interest in flux space (Fig. 3). Flux control, e.g. flux balance analysis (FBA) that focuses on steady state fluxes, has dominated recent progress on computational models of large metabolic networks and achieved significant advancement in both academic research and industrial applications [7], [8].

**Fig. 2:**
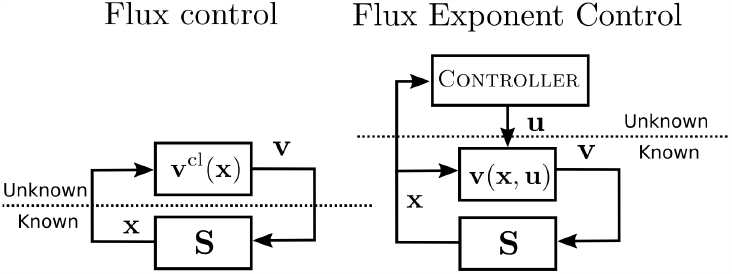
Control diagram comparison between flux control and FEC, with known-unknown splits depicted.

**Fig. 3:**
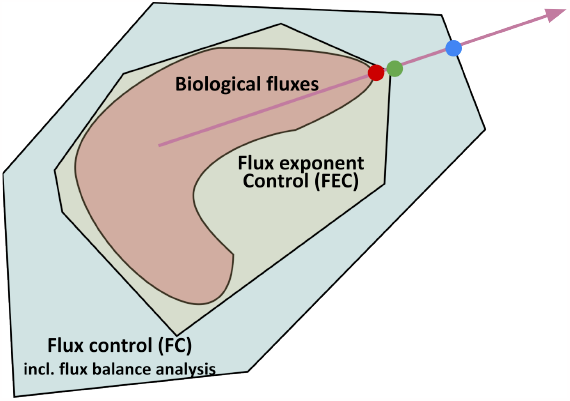
Illustration of constraint-based methods. The arrow represents the optimization objective, e.g., growth. The red set describes the actual set of biologically feasible actions that a cell can take. The red dot represents the biologically feasible action that optimizes the objective. The light blue outer-most set is the set of actions allowed by flux control, which includes flux balance analysis (FBA). It is only constrained by stoichiometry, therefore includes biological actions as a strict subset. The blue dot is the optimal action expected by flux control. The light green set denotes the set of actions constrained by flux exponent control (FEC), a strict super set of biological actions and a strict subset of flux control. The green dot is the optimal control action predicted by FEC.

However, flux control has difficulty capturing dynamic features of metabolic regulation. Not only do most flux control methods such as FBA assume metabolic fluxes are fast and therefore static (at steady state), flux control is also fundamentally unfit to model dynamics that are intrinsic to metabolic regulations. This is because the plant-controller split in flux control yields a plant with trivial dynamics (Fig. 1b and Sec. II-C). This is far away from reality, since it is known that metabolism in cells have strong plant dynamics, or dynamics without active regulation, that imposes significant limitations on control performance [9]. In other words, rather than controlling static fluxes in response to slow varying external environments on the timescale of days, metabolic control in cells may be dominated by concerns of a dynamic nature, such as stability, lag, oscillations, and growth arrest on the timescale of seconds to hours [10], [11]. From another perspective, since the strength of a constraint-based method comes from the set of constraints it could use, flux control is under-constrained and allows too many un-biological actions because its only constraint is the stoichiometry (see Fig. 3). As a result, predictions of flux control methods are often far from reality, unless extensive experimental data and expert knowledge are incorporated into the model via manual curation [12].

In this work, to resolve the above difficulties of flux control, we propose a novel constraint-based method called flux exponent control (FEC) to model metabolic dynamics as optimal control problems. FEC captures intrinsic metabolism dynamics via a novel plant-controller split between enzymes catalyzing metabolic reactions and the binding reactions regulating the enzymes’ activity (Sec. II). The resulting optimal control problems in FEC can be numerically solved via model predictive control. This is illustrated via the example of glycolysis, where oscillations that arise due to an unstable zero in the plant can be captured in FEC but not in flux control (Sec. III). In general, FEC brings metabolism dynamics to the realm of control system analysis and design.

## II. Constraint-based approaches from the layered architecture of metabolism

### A. Layered architecture of metabolism

Constraint-based modeling splits the mechanisms of a metabolic system into two parts: a slow-varying known part of which we have solid knowledge, and a fast-varying unknown part of which we have little knowledge. Then the known part is taken as constraints, and the unknown part is allowed to vary freely. The set of all feasible behaviors of the system are then behaviors achieved by varying the unknown parts, with the known parts held fixed. Fundamentally, for this split to be effective conceptually and mathematically, the system of concern needs to have a natural layered architecture so that the lower layer, the layer that already exists and is to be controlled, is known and fixed, and the higher layer, the layer controlling the lower layer, is unknown and varies. This split requires both a time-scale separation of dynamics in each layer, and a corresponding structural split in the organization of the system. Such layered architectures may be generally viewed as a result of adaptation to achieve optimal performance at diverse timescales using components that are limited by severe tradeoffs such as the speed-accuracy tradeoff [13], [14].

For our purpose of modeling metabolic dynamics, we can describe the layered architecture of metabolism as roughly consisting of three layers: the bottom metabolic stoichiometry layer, the middle enzyme regulation layer, and the top gene expression layer (Fig. 4). Metabolites are produced and degraded according to the reaction stoichiometry of the bottom layer, but with fluxes regulated by interactions between enzymes and regulatory metabolites and proteins of the middle layer, while the enzymes and proteins in turn are produced and degraded at the top layer. We introduce them sequentially below in concert with our formulation of metabolic dynamics.

**Fig. 4:**
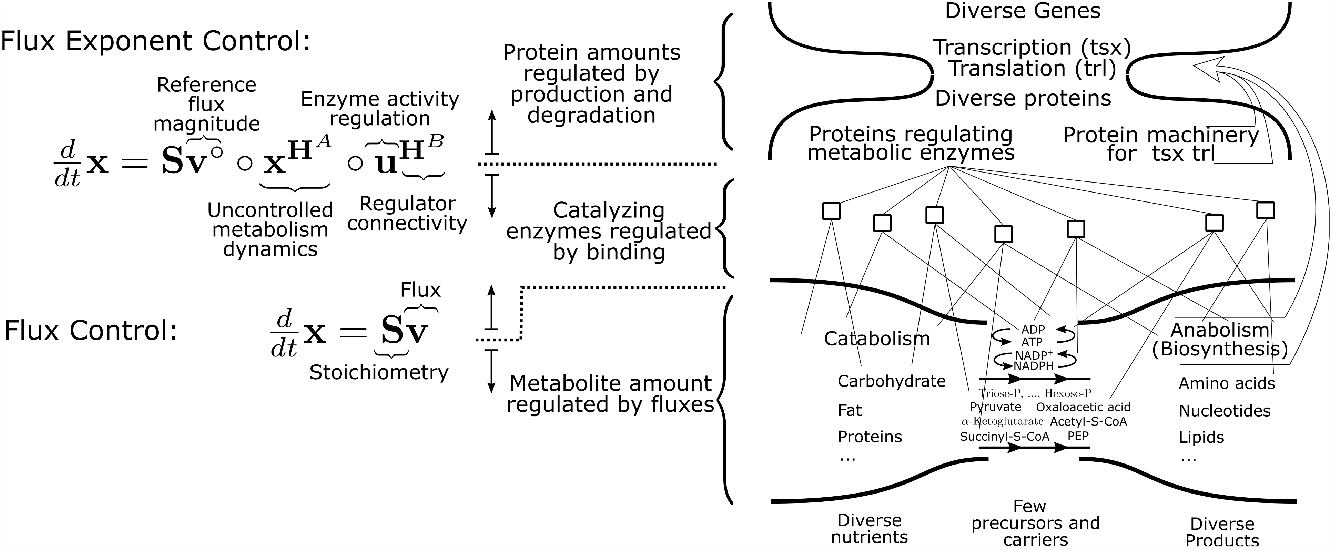
Illustration of how the two constraint-based approaches, flux control and flux exponent control, relate to splits in the layered architecture of metabolism. For simplicity, the terms for external exchange fluxes ***w*** are omitted. The layered architecture of metabolism is depicted on the right. Metabolism can be roughly viewed as consisting of three layers. The first (bottom) layer is the stoichiometry of metabolic reactions, capturing how metabolite amounts are regulated by reaction fluxes. The middle layer is the proteins’ regulation of metabolic enzymes, capturing how reaction fluxes are regulated by protein binding (squares). The top layer is transcription-translation, capturing how protein concentrations are regulated by production and degradation. This layer has an hourglass shape, connecting diverse genes with diverse proteins via a thin waist of transcription and translation machinery, with building blocks such as amino acids supplied by the bottom layer.

Viewed from below the bottom layer, we just have metabolites flowing in and out of the system. Without including any further structure from the layered architecture, the metabolite dynamics is a generic nonlinear dynamical system,

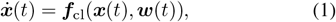

where 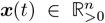 is the concentration of metabolite in the cell, 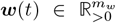 is exchange fluxes with external environments, and 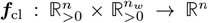 describes the change in metabolite concentrations. Here ***f***_cl_ is the closed loop dynamics, with the cell’s control actions on metabolism from all the layers of metabolic regulation included.

To have a more useful formulation of metabolic dynamics, we move up the layered architecture to include the bottom layer (Fig. 4). The bottom layer captures the knowledge that variations in metabolite concentrations are due to metabolic reactions, with the stoichiometry of these reactions describing the number of metabolite molecules consumed and produced. The reaction fluxes involved are catalyzed by enzymes with rates at millisecond to second timescales, much faster than any possible change to the metabolism stoichiometry. Therefore the stoichiometry can be considered as the fixed structure of the bottom layer, with quickly varying fluxes regulated by higher layers.

Incorporating the structure of the bottom layer therefore yields the *stoichiometry-flux split*, so we can write

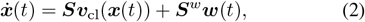

where ***S*** ∈ ℝ^*n*×*m*^ is the stoichiometry matrix of the cell’s internal metabolic reactions, *m* is the number of internal reactions, and 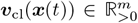 is the (closed loop) fluxes of these reactions, varying with metabolite concentrations through regulatory mechanisms such as enzyme allostery and gene regulation described in higher layers. 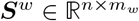 is the stoichiometry for the external exchange fluxes ***w***(*t*).

### B. Flux control from stoichiometry-flux split

The stoichiometry-flux split yields a constraint-based formulation called flux control, with the stoichiometry as constraints and the fluxes as control variables (Fig. 1). Indeed, since ***v***_cl_ represents the flux regulated by all the higher layers, if we do not have any knowledge about the structure of the higher layers, then ***v***_cl_ should be left as arbitrary. Therefore, a generic flux control formulation is the following:

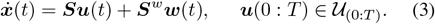

Here 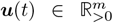 is the control variable, representing metabolic fluxes that can be controlled by the higher layers. Since the fluxes can vary with time, ***u***(0 : *T*) denotes the time trajectory of fluxes in the time interval [0, *T*] and 𝒰_(0:*T*)_ denotes the constraint on the flux trajectories, e.g., upper bounds on flux magnitudes from maximum enzyme catalysis rates and enzyme amounts in cells.

While constraints on flux trajectories can be hard to come up with, it is easier to constrain steady state fluxes. One popular flux control method, *flux balance analysis (FBA)*, does this by assuming steady states exist and are achieved. This assumption is applicable whenever the phenomenon of concern is much slower than cell metabolism, e.g., tens of hours or longer. Restricted to steady state fluxes, (3) becomes

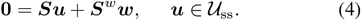

Here 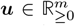 is the steady state fluxes, ***w*** is steady state exchange fluxes, and the constraint set has become static as well, 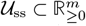. Further constraints on the static fluxes can come from thermodynamics of the reactions.

### C. Flux control ignores intrinsic metabolic dynamics

Flux control has one severe limitation: the lack of internal dynamics. We can understand this clearly by viewing the constraint-based formulation that turn stoichiometry-flux split (2) into flux control (3) as using the layered architecture of metabolism to formulate a control system. Without the knowledge of the bottom layer, we have cellular metabolism as the following generic control system:

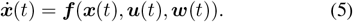

If we know the control law ***u*** of how metabolic fluxes are controlled by the cell, e.g., a static function ***u***(***x***), we can close the loop to yield ***f***_cl_(***x, w***) = ***f***(***x, u***(***x***), ***w***) as in (1).

Since the stoichiometry-flux split (2) does not incorporate knowledge of how the fluxes are regulated in the higher layers, we are forced to take the fluxes as control variables, therefore obtaining the flux control formulation (3) (Fig. 2). Because the stoichiometry simply multiplies the fluxes to yield metabolite concentration dynamics, the resulting plant-controller split has trivial plant dynamics (Fig. 1). If we want to incorporate nontrivial plant dynamics at the flux level, we need to specify the full dynamics of fluxes, i.e., the exact form of the function ***v***_cl_(***x***), defeating the purpose we started with: to model metabolic dynamics without detailed knowledge of flux dynamics.

Therefore, to capture intrinsic metabolic dynamics, we need knowledge from the higher layers about how fluxes are regulated. This brings the middle layer to our attention, which states that metabolic fluxes are catalyzed by enzymes, and enzymes’ activities are regulated to change fluxes (Fig. 4). One implication of this is that while the fluxes are regulated, they also have un-regulated dynamics since the enzymes continue to catalyze metabolites and produce metabolic fluxes without regulations of enzymes’ activities. So instead of the closed loop flux ***v***_cl_(***x***) in (2) that only depends on ***x***, or ***v*** = ***u*** in flux control (3) where all of fluxes are controlled, we write ***v***(***x, u***) and

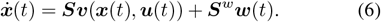

The control variable ***u*** here are regulatory actions on the fluxes from the middle and higher layers (Fig. 2). To make (6) useful and formulated into a constraint-based approach, we need to characterize how ***v*** depends on ***x*** and ***u*** to refine the functional form of ***v***(***x, u***). This structure of how fluxes are regulated is described in the middle layer, which we analyze in detail next.

### D. Flux exponent control (FEC) from binding-catalysis split

Above the bottom layer of metabolic reactions, the middle layer specifies that metabolic fluxes are catalyzed by enzymes, which in turn are regulated by the binding with metabolites, cofactors, and proteins, or transformation of molecular states by covalent modifications such as phosphorylation or methylation (Fig. 4). The binding reactions in the middle layer regulating enzyme activities are naturally separated from the bottom and top layers in time scales [15], [16]. These binding reactions often reach equilibrium on a millisecond to second timescale, while changes in metabolite concentrations of the bottom layer take tens of seconds or longer, and production and degradation of proteins in the top layer take tens of minutes to hours. This timescale separation yields a split between catalytic enzymes and their binding regulations, or a *binding-catalysis split*.

To formulate the binding-catalysis split mathematically, we need to parameterize the set of control actions on catalysis fluxes allowed by binding reactions. As described in Chapter 2 and 3 of [17] (also [18]), the full regulatory profile of enzyme activities via binding reactions can be parameterized as adjusting the reaction orders, or exponents, of ***v***’s dependence on ***x***, constrained in a polyhedral set.

This means, the reaction order, mathematically defined as the log derivative, of the closed-loop flux ***v***_cl_ to metabolite concentrations ***x***, denoted ***H***^cl^ with 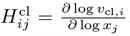, is the open-loop gain ***H***^*A*^, or passive reaction orders of fluxes to metabolites, describes how the reactions would proceed without regulations of the catalyzing enzymes by binding closed loop gain under the regulation of binding reactions. This reaction order ***H***^cl^, or closed-loop gain, is further constrained in a polyhedral set that is directly specified by the binding reaction network’s stoichiometry. Just like metabolic network stoichiometry is the structure of the bottom layer, the binding network stoichiometry is the structure of the middle layer. With this characterization, the binding-catalysis split is mathematically a *rule* of bioregulation that cells regulate metabolic fluxes by adjusting their exponents, or reaction orders. We call this rule *flux exponent control* (FEC).

To formulate FEC into a constraint-based approach, we need to turn the closed-loop description of fluxes in (2) into an open-loop or control system description in (6). This means, instead of the closed loop flux ***v***_cl_(***x***) that includes all upper layers’ regulations, we have open loop flux ***v***(***x, u***), with ***u*** the control variable tuned by metabolites and protein concentrations in layers other than the middle. The reaction orders, or gains, of the open loop and closed loop can be defined locally around an operating point (***x***_0_, ***u***_0_), with 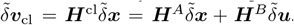. Here 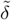 denotes fold-change variations, e.g., 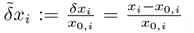, 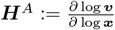 is the open-loop gain of fluxes to metabolite concentrations, and 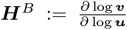 is the connectivity of which fluxes are regulated by which control variables, with log applied component-wise. If we close the loop with a static controller map ***u***(***x***) and define 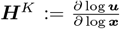, then ***H***^cl^ = ***H***^*A*^+***H***^*B*^***H***^*K*^, analogous to the static state feedback case in linear systems. Therefore, polyhedral constraints on closed-loop gain ***H***^cl^ from binding network stoichiometry propagates to polyhedral constraints on controller gain ***H***^*K*^.

Now with a clear understanding of the local description of FEC, we can integrate to obtain a global description. The open loop gain ***H***^*A*^,or passive reaction orders of fluxes to metabolites, describes how the reactions would proceed without regulations of the catalyzing enzymes by binding reactions. Therefore, we can often consider ***H***^*A*^ as the mass-action reaction orders, so that ***H***^*A*^ is constant. ***H***^*B*^ is also constant since it simply captures the connectivity of fluxes’ regulation. With constant ***H***^*A*^ and ***H***^*B*^, we can integrate from an operating point (***x***_0_, ***u***_0_) to obtain log ***v*** = ***H***^*A*^ log ***x*** + ***H***^*B*^ log ***u*** + ***c*** for some constant vector ***c***. This ***c*** can be considered as log ***v*** at some standard values of ***x*** and ***u***, denoted ***x***° and ***u***°. So we define the standard reference flux log ***v***° := ***c*** = log ***v***(***x***°, ***u***°). Note that the standard reference flux ***v***° simply defines the “units”, with no relation to the flux at an operating point ***v***_0_ = ***v***(***x***_0_, ***u***_0_), which can be arbitrarily chosen. Now we can write,

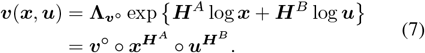

Here exponential is applied component-wise to a vector. The operation ○ denotes Hadamard or element-wise product. 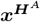 denotes the vector with 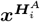 as the *i*th element, where we use the notation 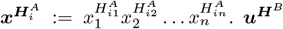 is similarly defined.

With the FEC structure of the middle layer on how the fluxes are regulated in (7), we can incorporate it into the stoichiometry-flux structure of the bottom layer (6) to have the FEC formulation of metabolism as a control system:

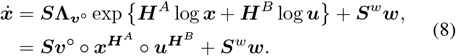

## III. Solving FEC using model predictive control

As a constraint-based approach, FEC parameterizes the space of all feasible behaviors in a metabolic system by formulating dynamic regulations as controller designs. This way, FEC can use control theory tools to derive hard limits on performance for all possible controllers from metabolic system structures, as in [9]. Beyond general laws, we are also interested in solving for metabolic dynamics in specific scenarios. Since constraint-based approaches already define the set of feasible control actions, we can solve for a specific scenario by formulating an optimization problem that search for relevant actions in this set. We just need to specify the scenario via an optimization objective and some additional constraints. For FBA (4), this is often a linear or quadratic program that yields a vector of steady state fluxes [7]. For the FEC formulation of metabolism (8), this is an optimal control problem that yields a dynamic trajectory of metabolite concentrations ***x***, control of flux exponents ***u***, and fluxes ***v***. This nonlinear optimal control problem with state and controller constraints can then be solved using model predictive control (MPC) methods [19], [20]. The generic optimal control problem of FEC is the following:

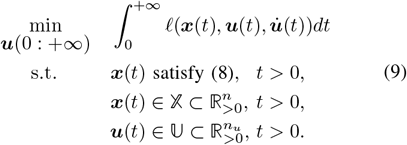

Here the time horizon is considered infinite, and the objective function is the integral of a per-time loss *ℓ*(·) over this infinite time horizon. Loss *ℓ* can contain the linear-quadratic cost relative to reference values to encode the objective of maintaining a steady state. It can contain the negation of a sum of metabolite concentrations ***x*** to promote yield maximization of biomass or certain metabolites. It can also contain quadratic cost on 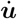 to penalize regulation of fluxes by changing concentrations of molecules. The variable to be minimized over is the control action trajectory ***u***(0 : +). The first constraint is the dynamic equation from FEC (8). The second constraint requires that states, or metabolite concentrations ***x***, are contained in the set 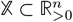 for all time. This can include lower bounds on ATP concentration for example. The third constraint requires that control variables ***u*** are in 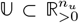 for all time. This can include actuator saturation from bounds on flux magnitudes or reaction orders(Sec. II-D), for example.

To solve (9) computationally, we use the MPC formulation to consider a local problem at a given time *t* with state (***x***(*t*), ***u***(*t*), ***w***(*t*)) [19], [20]. We linearize the system around this point, and consider a local optimal control problem of this linearized system with *T* time horizon with discrete time step ∆*t*. The solved optimal controller is then implemented for one step in ***u***(*t*′) for *t*′ = *t* to *t* + ∆*t*. Then we consider a local problem at time *t* + ∆*t* and solve again.

### A. Example: glycolytic oscillation

We illustrate the capability for FEC to capture metabolic dynamics via the example of oscillations in glycolysis. Glycolytic oscillations are well studied biologically [21] and from a control theory perspective [9]. It is known to happen on the timescale of 30 seconds, much shorter than the timescale for metabolism to reach steady state [21]. It is understood that the system oscillates due to aggressive control actions, implemented by allosteric regulation of enzymes, that adapts to changing supplies and demands. This attenuation of steady state error causes oscillation when hit by large disturbances, made more severe by the autocatalytic stoichiometry that is intrinsically unstable. FEC promises that just based on the stoichiometry of this system, we have a parameterization of all possible regulations the cell can take on glycolysis. Then by simply asking for controllers that aggressively attenuate steady state error, we should be able to uncover oscillatory response, with no information other than the stoichiometry. We demonstrate this below.

Following [9], instead of considering the detailed steps of reactions in glycolysis, we consider two lumped reactions that capture the structure of autocatalysis. This yields the following dynamics for the concentrations of metabolites:

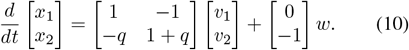

Here *x*_2_ is ATP, or energy charge, and *x*_1_ is a lumped intermediate of the glycolysis pathway, such as fructose 1,6bisphosphate. The first reaction with flux *v*_1_ consumes *q* units of ATP and produce one unit of intermediate. The second reaction with flux *v*_2_ consumes one unit of intermediate and produces 1 + *q* units of ATP. Together, looping through the two reactions once results in a net production of one unit of ATP. We also include an external disturbance with flux *w* that consumes one unit of ATP. This corresponds to the maintenance energy cost of the cell, which can increase under environmental disturbances such as heat shocks.

To formulate into FEC, we write 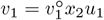, and 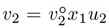, as described in (7) in Sec. II-B. Here *v*_1_ has unregulated dependence on *x*_2_ because of mass action and that *x*_2_, ATP, is a reactant of *v*_1_. Similarly, *v*_2_ has un-regulated dependence on *x*_1_ because *x*_1_, the intermediate, is a reactant for the second reaction. For the loss function *ℓ*(·), we use the following linear quadratic cost in fold-change variables:

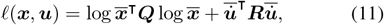

where 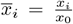 is the fold-change difference between the current state and the desired state, 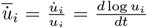 is the fold-change time derivative of ***u***, and we add a lower bound on the ATP level as a state constraint, *x*_2_(*t*) ≥ 0.6. Time derivative of ***u***, not ***u***, is penalized since without changing molecular concentrations in binding regulations, the fluxes stay constant rather than going back to a reference value.

From the simulation result in Fig. 5, we see FBA (orange line) cannot capture the oscillations, while FEC can, directly from the autocatalytic stoichiometry of glycolysis. FEC also captures the speed-robustness tradeoff, the central feature of this system [9], that more aggressive feedback regulation causes better steady state adaptation but larger oscillations.

**Fig. 5:**
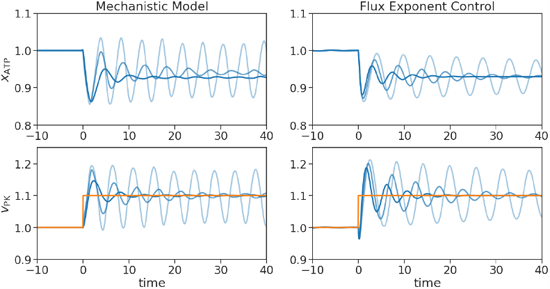
Simulations for glycolytic oscillation via the mechanistic model in [9] (left) or via FEC solved by MPC (right). The steady state fluxes predicted by FBA is the orange line. *x*_ATP_, or *x*_2_ in the text, is concentration of ATP. *v*_PK_, or *v*_2_ in the text, is the reaction flux consuming intermediate and producing ATP. The parameter values of the mechanistic model, in notations of [9], are *a* = 1, *q* = 1, *k* = 1.1, and *g* = 0.3. *h* = 2.5, 2.8, or 3.1, for the trajectories from dark to light blue, with increasing oscillation magnitude. The disturbance *w* is 1 from *t* = − 10 to 0, and jumps to 1.1 for *t >* 0. Parameters in the formulation of FEC are chosen to match with steady state values of the mechanistic model: 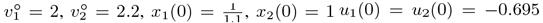 and ***x***_0_ = ***x***(0) when *w* = 1, then (0.93, 0.98) when *w* = 1.1. The cost and optimization parameters of FEC are ∆*t* = 0.03, *T* = 0.6, ***Q*** = diag(0.3, 0.08) and ***R*** (in the local problem in discrete time) is diag(3, 3), diag(4.25, 4.25) or diag(5.5, 5.5), for the three trajectories from dark to light blue with increasing oscillation magnitudes. Code is available at https://github.com/chemaoxfz/FEC-ACC-202303

## IV. Discussion

In this work, we reveal a general link between constraint-based modelling methods and plant-controller splits in the layered architecture of metabolism. This clarifies the fundamental limitation of flux control, including FBA, in capturing metabolism dynamics. Overcoming this limitation, we develop FEC based on a structural split between intrinsic dynamics of metabolic fluxes and their regulations. As a constraint-based approach, FEC formulates metabolism as a control system, with metabolic dynamics solvable as optimal controllers via MPC. FEC can capture metabolic dynamics at seconds to hours timescale, such as glycolytic oscillations. Therefore, FEC has great potential in applications from metabolic engineering of microbial communities to understanding cell growth and survival in dynamic environments.

To apply FEC to compute metabolic dynamics for largescale problems such as whole-genome models in application, we need to solve the optimal control problem in (9) at scale. This can be achieved by combining FEC with system level synthesis [22] so that distributed and localized MPC methods [23] can utilize the network sparsity and controller locality in metabolism to speed up computation.

More generally, FEC opens the door to analysis and design that map metabolic system architecture to dynamic functions. This may have far-reaching implications on drug targets and engineering of synthetic organisms. For example, if instead of an antibiotic targeting a protein, we design treatments targeting hard limits, e.g., speed-robustness tradeoffs, due to regulatory architectures of microbes. Then since it is hard to change the architecture via evolution that accumulates greedy small steps, such architecture-targeted treatments would be hard to escape through mutation. As another example, instead of inserting a gene to make a microbe fit in a static environment, we can engineer a regulatory architecture in the microbe to adapt to a dynamic environment with large shifts. Then this microbe could gain a dynamic niche in the time domain, and persistently survive in an environment without requiring an overall growth advantage. This may be a promising path to establish an engineered microbial species in an environment of deployment in applications [4].

## References

[1] D. W. Erickson, et al., “A global resource allocation strategy governs growth transition kinetics of escherichia coli,” Nature, vol. 551, no. 7678, pp. 119–123, 2017.

[2] T. Khazaei, et al., “Metabolic multistability and hysteresis in a model aerobe-anaerobe microbiome community,” Science Advances, vol. 6, no. 33, 2020.

[3] R. Blumberg and F. Powrie, “Microbiota, disease, and back to health: a metastable journey,” Science translational medicine, vol. 4, no. 137, pp. 137rv7–137rv7, 2012.

[4] M. B. N. Albright, et al., “Solutions in microbiome engineering: prioritizing barriers to organism establishment,” The ISME Journal, vol. 16, no. 2, pp. 331–338, Feb 2022.

[5] S. R. Lindemann, et al., “Engineering microbial consortia for controllable outputs,” The ISME Journal, vol. 10, no. 9, pp. 2077–2084, Sep 2016.

[6] P. d. R. Martins Conde, T. Sauter, and T. Pfau, “Constraint based modeling going multicellular,” Frontiers in Molecular Biosciences, vol. 3, 2016.

[7] B. Ø. Palsson, Systems Biology: Constraint-based Reconstruction and Analysis, stu - student edition ed. Cambridge University Press, 2015.

[8] L. Heirendt, et al., “Creation and analysis of biochemical constraint-based models using the cobra toolbox v.3.0,” Nature Protocols, vol. 14, no. 3, pp. 639–702, Mar 2019.

[9] F. A. Chandra, G. Buzi, and J. C. Doyle, “Glycolytic oscillations and limits on robust efficiency,” Science, vol. 333, no. 6039, pp. 187–192, 2011.

[10] E. de Alteriis, et al., “Revisiting the crabtree/warburg effect in a dynamic perspective: a fitness advantage against sugar-induced cell death,” Cell Cycle, vol. 17, no. 6, pp. 688–701, 2018, pMID: 29509056.

[11] S. Devkota, et al., “Dietary-fat-induced taurocholic acid promotes pathobiont expansion and colitis in il10-/-mice,” Nature, vol. 487, no. 7405, pp. 104–108, Jul 2012.

[12] L. Coppens, et al., “Vibrio natriegens genome-scale modeling reveals insights into halophilic adaptations and resource allocation,” Molecular Systems Biology, vol. n/a, no. n/a, p. e10523.

[13] J. C. Doyle and M. Csete, “Architecture, constraints, and behavior,” Proceedings of the National Academy of Sciences, vol. 108, no. Supplement 3, pp. 15 624–15 630, 2011.

[14] Y. Nakahira, et al., “Diversity-enabled sweet spots in layered architectures and speed accuracy trade-offs in sensorimotor control,” Proceedings of the National Academy of Sciences, vol. 118, no. 22, p. e1916367118, 2021.

[15] G. E. Briggs and J. B. S. Haldane, “A Note on the Kinetics of Enzyme Action,” Biochemical Journal, vol. 19, no. 2, pp. 338–339, 01 1925.

[16] J. Gunawardena, “A linear framework for time-scale separation in nonlinear biochemical systems,” PLOS ONE, vol. 7, no. 5, pp. 1–14, 05 2012.

[17] F. Xiao, “Biocontrol of biomolecular systems: Polyhedral constraints on binding’s regulation of catalysis from biocircuits to metabolism,” Ph.D. dissertation, California Institute of Technology, 2022.

[18] F. Xiao, M. Khammash, and J. C. Doyle, “Stability and control of biomolecular circuits through structure,” in 2021 American Control Conference (ACC), 2021, pp. 476–483.

[19] D. Q. Mayne, “Model predictive control: Recent developments and future promise,” Automatica, vol. 50, no. 12, pp. 2967–2986, 2014.

[20] F. Borrelli, A. Bemporad, and M. Morari, Predictive Control for Linear and Hybrid Systems. Cambridge University Press, 2017.

[21] S. Danø, P. G. Sørensen, and F. Hynne, “Sustained oscillations in living cells,” Nature, vol. 402, no. 6759, pp. 320–322, Nov 1999.

[22] J. Anderson, et al., “System level synthesis,” Annual Reviews in Control, vol. 47, pp. 364–393, 2019.

[23] C. A. Alonso, et al., “Distributed and localized model predictive control. part i: Synthesis and implementation,” IEEE Transactions on Control of Network Systems, pp. 1–12, 2022.

